# The role of control region mitochondrial DNA mutations in cardiovascular disease: stroke and myocardial infarction

**DOI:** 10.1101/382374

**Authors:** Miriam Umbria, Amanda Ramos, Maria Pilar Aluja, Cristina Santos

## Abstract

Recent studies associated certain type of cardiovascular disease (CVD) with specific mitochondrial DNA (mtDNA) defects, mainly driven by the central role of mitochondria in cellular metabolism. Considering the importance of the control region (CR) on the regulation of the mtDNA gene expression, the aim of the present study was to investigate the role of the mtDNA CR mutations in two CVDs: stroke and myocardial infarction (MI). Both, fixed and heteroplasmy mutations of the mtDNA CR in two population samples of demographically-matched case and controls, were analysed using 154 stroke cases, 211 MI cases and their corresponding control individuals. Significant differences were found between cases and controls, reporting the m.16145G>A and m.16311T>C as a potential genetic risk factors for stroke (conditional logistic regression: p=0.038 and p=0.018, respectively), whereas the m.72T>C, m.73A>G and m.16356T>C could act as possible beneficial genetic factors for MI (conditional logistic regression: p=0.001, p=0.009 and p=0.016, respectively). Furthermore, our findings also showed a high percentage of point heteroplasmy in MI controls (logistic regression: p=0.046; OR= 0.209, 95% CI [0.045-0.972]). These results demonstrate the possible role of mtDNA mutations in the CR on the pathogenesis of stroke and MI, and show the importance of including this regulatory region in genetic association studies.

**Author Summary:** Given the association between cardiovascular disease and specific mitochondrial DNA (mtDNA) defects and considering the importance of the control region of this genome on the regulation of mtDNA gene expression, here, we investigate the role of mutations in mitochondrial DNA control region in two cardiovascular diseases: stroke and myocardial infarction. In this study we found five mitochondrial genetic variants related to cardiovascular disease, based on single nucleotide polymorphisms (SNPs), which are located in the control region of mtDNA. Despite the abundance of work on the role of mitochondrial DNA in relation to cardiovascular disease, little literature has been published on the variation that this genome expresses in relation to this disease. For this reason, our study provides significant insight of the genetic variability that determines normality or pathology in relation to the genetic risk of cardiovascular disease. The results obtained demonstrate the possible role of mtDNA mutations in the control region on the pathogenesis of stroke and myocardial infarction, and show the importance of including this regulatory region in genetic association studies.

## Introduction

Cardiovascular disease (CVD) is one of the most widespread and common causes of death in the world. The onset and severity of these diseases are influenced by both genetic and environmental factors. Recent evidences associate mitochondrial dysfunction with several cardiovascular manifestations, mainly driven by the central role of mitochondria in cellular metabolism, particularly in energetically demanding tissues such as the brain and heart [1,2].

Human mitochondrial DNA (mtDNA) is 16.6-kb double-stranded circular DNA molecule that encodes for 13 electron transport chain (ETC) proteins, 2 ribosomal RNAs (rRNAs) and 22 transports RNAs (tRNAs). The control region (CR) encompasses the light and heavy strand promoters, the heavy strand origin of replication (O_H_), three conserved sequence blocks and the termination associated sequences (TAS) [3]. MtDNA is more susceptible than nuclear DNA to oxidative damage, probably due to the lack histone complex and an inefficient DNA repair mechanisms, which may serve as a protective barrier against external and internal noxious agents as reactive oxygen species (ROS) [4]. However, the hypothesis of direct damage by ROS is increasingly criticized and it is suggested that errors in mtDNA replication and repair may be the main cause of its high mutation rate (~10-fold greater than in nDNA) [5].

Recent evidence have linked certain CVDs with specific mtDNA mutation including base substitution [6–11], deletions [12], duplications [13] and point or length heteroplasmy [14–17] both in coding [6,9,10,12,14,15] and noncoding region [6–9,11,13,16,17] of mtDNA. In particular, mtDNA mutations located in CR have a potential importance since they may influence on the regulation of the mtDNA gene expression. In fact, several studies detected association of a great range of mtDNA variants (with negative or beneficial effect, both fixed or in heteroplasmy) and different diseases [6,9].

In general, these disparities could occur because mtDNA mutations in the CR may not be directly tied to any form of pathology, but could capable of influencing mitochondrial function through changes in the number of copies, inducing profound effects on the expression of mitochondrial-encoded gene transcripts and related enzymatic activities (complexes I, III, and IV) [18,19].

The ability of mtDNA mutation to influence in the development of CVDs is directly related to its prevalence and the severity of its impact on mitochondrial function. In addition, several studies have demonstrated that due to the differences in the prevalence of the main etiological factors between intra- and extracranial arteries, the effect of the mtDNA mutations with stroke and MI could be different [20–23]. The main aim of the present study was to investigate the role of CR mtDNA mutations (fixed or in heteroplasmy) in two CVDs; stroke and MI.

## Results

### Analysis of fixed and heteroplasmic mtDNA mutations with stroke

A detailed matrix of all mtDNA positions analyzed in stroke cases and controls are reported in S2 Table and the frequencies of fixed mutations found are showed in Table 1. The percentages of m.16145G>A and m.16311T>C were overrepresented in stroke cases (5.2% and 18.2%, respectively) than controls (1.9% and 9.7%, respectively). After correction for the effect of CV risk factors with significant differences between stroke cases and controls (hypercholesterolemia [24]), significant association were still observed in m.16145G>A (conditional logistic regression: p=0.038; OR= 4.407, 95% CI [1.086-17.883]) and m.16311T>C (conditional logistic regression: p=0.018; OR= 2.417, 95% CI [1.165-5.016]), emerging as a possible genetic risks factors for stroke (Table 1).

**Table 1:**
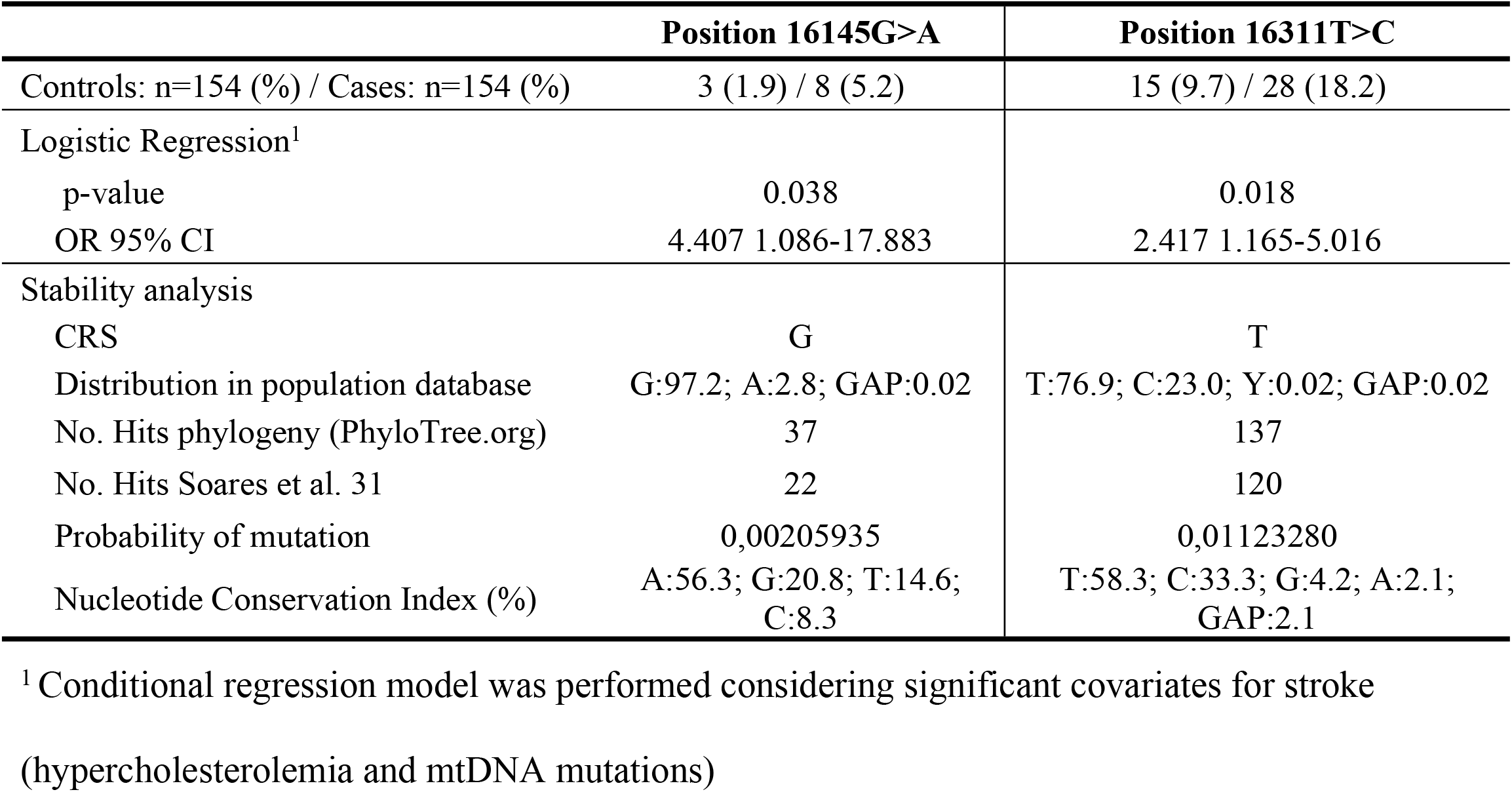
Complete results of stroke fixed mtDNA mutation analysed

|  | Position 16145G>A | Position 16311T>C |
| --- | --- | --- |
| Controls: n=154 (%) / Cases: n=154 (%) | 3 (1.9) / 8 (5.2) | 15 (9.7) / 28 (18.2) |
| Logistic Regression<sup>1</sup> |  |  |
| p-value | 0.038 | 0.018 |
| OR 95% CI | 4.407 1.086-17.883 | 2.417 1.165-5.016 |
| Stability analysis |  |  |
| CRS | G | T |
| Distribution in population database | G:97.2; A:2.8; GAP:0.02 | T:76.9; C:23.0; Y:0.02; GAP:0.02 |
| No. Hits phylogeny (PhyloTree.org) | 37 | 137 |
| No. Hits Soares et al. 31 | 22 | 120 |
| Probability of mutation | 0,00205935 | 0,01123280 |
| Nucleotide Conservation Index (%) | A:56.3; G:20.8; T:14.6; C:8.3 | T:58.3; C:33.3; G:4.2; A:2.1; GAP:2.1 |

<sup>1</sup> Conditional regression model was performed considering significant covariates for stroke (hypercholesterolemia and mtDNA mutations)

Stability analyses were performed to predict the impact of these mutations. Several measures as the number of hits in the mtDNA phylogeny, the probability of mutation, the frequency in the population database and the conservation index (CI) at nucleotide level, were calculated, and results are showed in Table 1. The results obtained revealed m.16145G>A and m.16311T>C as non-stable position since they present a minimum of 37 hits in the phylogeny, a high probability of mutation, a high frequency of the variant in the population database (here denoted by minor allele frequency [MAF] >5%) detected on m.16311T>C or low-frequency (MAF 1-5%) in m.16145G>A and a maximum nucleotide CI of 58% (Table 1). To infer about the impact of m.16145G>A and m.16311T>C on the stability of secondary structures of the mtDNA, a prediction of different structures with the wild type (rCRS) and mutant variant was performed. It seems that m.16311T>C implies a conformational rearrangement, resulting in structure of Fig 1 as the new predicted minimum free energy solution (-0.40 kcal/mol), causing a stability reduction of the region. No structural or thermodynamic differences were found for the m.16145G>A.

**Fig 1.**
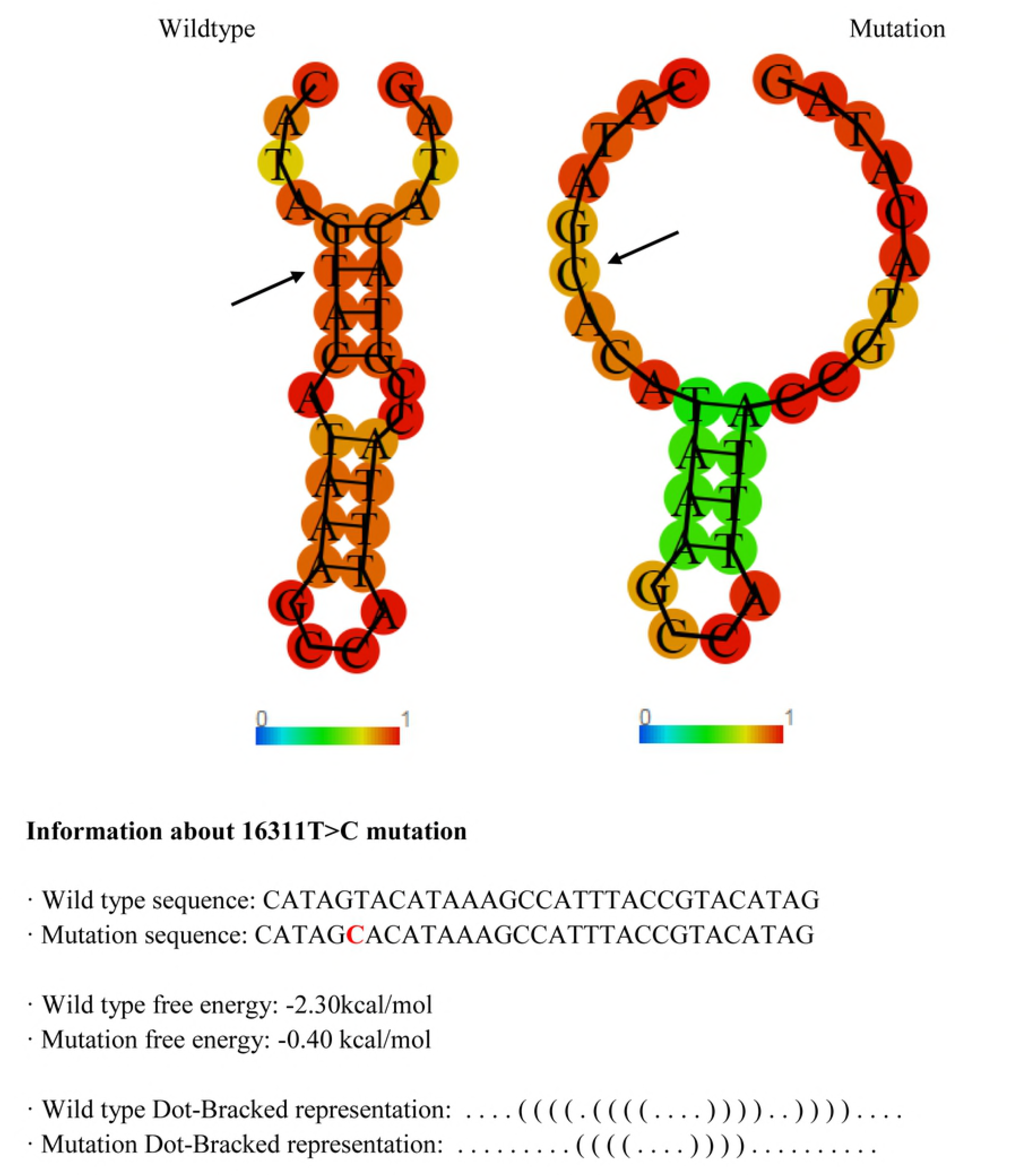
Wild type vs. mutant structure and energy information. For m.16311T>C, relevant secondary structure and energy information is listed along with a graphical drawing for both the mutant and the wild type.

The distribution of the heteroplasmic positions between stroke cases and controls are reported in Table 2. Eighty-eight stroke cases (57.1%) and eighty-six controls (55.8%) presented point and/or length heteroplasmy, and no significant differences were obtained between groups (PH: p=1.000 and LH: p=1.000, McNemar’s test). The most prevalent variant detected was a length heteroplasmy located in the poly-C tract of the HVRII (between positions 303–315 of the mtDNA), which was present in a 52% of stroke cases and in 46.7% of controls. Point heteroplasmies were found in six stroke cases and six controls, involving nine different positions of the mtDNA: 146, 150, 152, 185, 204, 16092, 16093, 16129 and 16399.

**Table 2:**
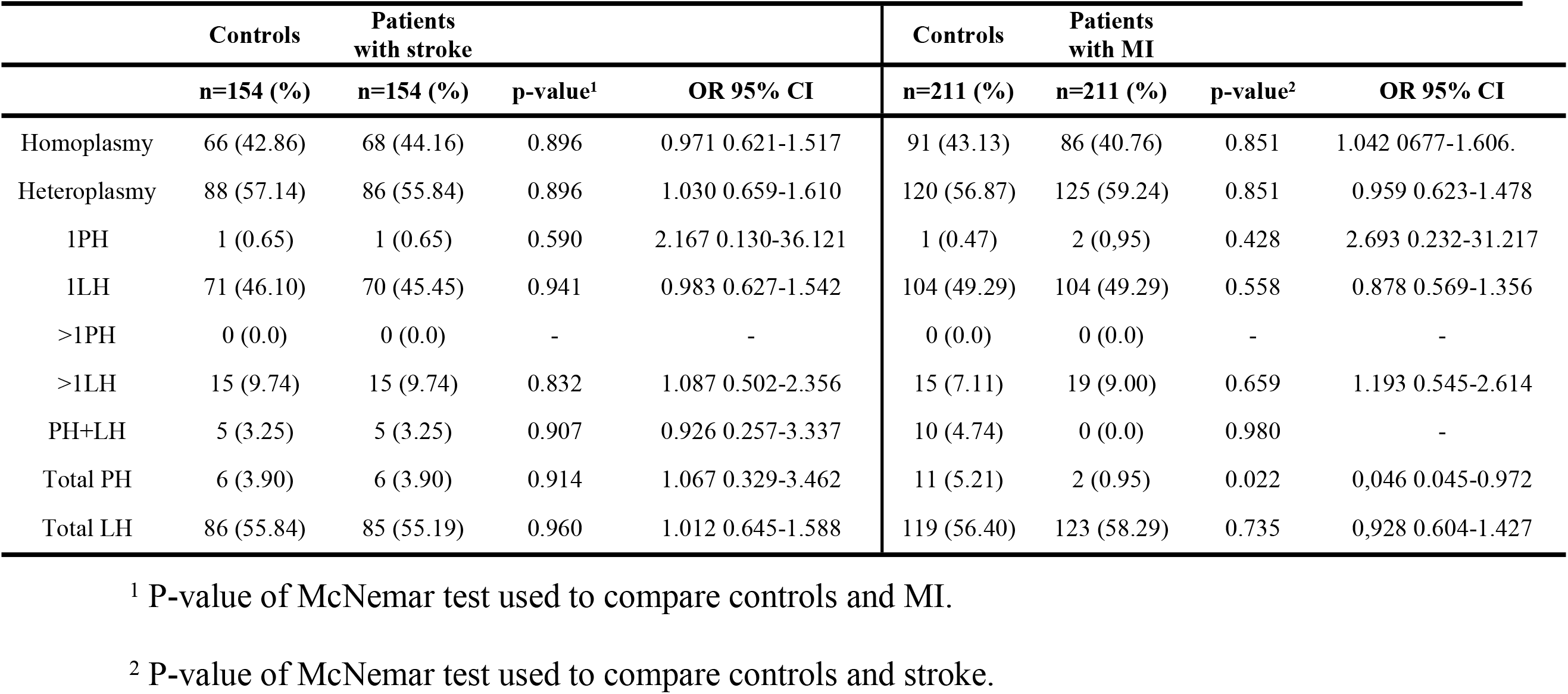
Classification of the analyzed stroke and myocardial infarction (MI) individuals depending on the type(s) of heteroplasmy they presented

The analysis of stability performed to predict the impact of these heteroplasmic positions is presented in Table 3. In general, these mutations have a minimum of 16 hits in the phylogeny, were located in hotspots positions, have a high frequency of the minor variant in the population database (min. height peaks 16.67%), and a low conservation index, indicated these heteroplasmies have typical characteristics of non-stable positions (Table 3).

**Table 3:** Complete results of stroke and myocardial infarction heteroplasmy position analyzed (position, sample name, heteroplasmy type, heteroplasmy origin, distribution in population database, number of hits in mtDNA phylogeny (PhyloTree.org) and by Soares *et al*,[25] probability of mutation and nucleotide Conservation Index)

| Position | Sample name | Het | CRS | Mean proportion height peaks | Distribution in population database | No. Hits phylogeny (PhyloTree.org) | No. Hits Soares et al. 31 | Probability of mutation | Nucleotide Conservation Index (%) |
| --- | --- | --- | --- | --- | --- | --- | --- | --- | --- |
| 146 | Ca10_Stroke | T/c | T | 83.33T 16.67C | T:81.4; C:18.4; A:0.1; GAP:0.1 | 121 | 109 | 0,01020313 | C:41.7; T:31.3; A:27.1 |
| 150 | Ca111_Stroke | T/c | C | 68T 32C | C:88.2; T:11.7; G:0.1; GAP:0.1 | 74 | 63 | 0,00589722 | C:43.8; G:27.1; A:18.8; T:10.4 |
| 152 | Ca69_Stroke | C/t | T | 52.94C 47.06T | T:70.4; C:29.5; GAP:0.1; G:0.02 | 196 | 157 | 0,01469625 | C:47.9; T:35.4; A:16.7 |
| 185 | Ca115_Stroke | G/a | G | 66.67G 33.33A | G:94.6; A: 3.9; T: 1.1; C:0.3; GAP: 0.1; R:0.02 | 24 | 24 | 0,00224656 | C:52.1; A:29.2; T:12.5; G:6.3 |
| 204 | Ca79_Stroke | T/c | T | 54.55T 45.45C | T:93.4, C:6.5, A:0.1, GAP:0.1; Y: 0,02 | 44 | 43 | 0,00402509 | G:52.1; T:22.9; C:16.7; A:8.3 |
| 16129 | Ca13_Stroke | G/a | A | 73.33G 26.67A | G:84.6; A: 14.9; C: 0.4; R:0.02; GAP: 0.02 | 93 | 86 | 0,00805017 | A:87.5; G:8.3; T:2.1; GAP:2.1 |
| 146 | Co07_Stroke | T/c | T | 69.23T 30.77C | T:81.4; C:18.4; A:0.1; GAP:0.1 | 121 | 109 | 0,01020313 | C:41.7; T:31.3; A:27.1 |
| 146 | Co68_Stroke | C/t | T | 80C 20T | T:81.4; C:18.4; A:0.1; GAP:0.1 | 121 | 109 | 0,01020313 | C:41.7; T:31.3; A:27.1 |
| 146 | Co35_Stroke | T/c | T | 66.67T 33.33C | T:81.4; C:18.4; A:0.1; GAP:0.2 | 121 | 109 | 0,01020313 | C:41.7; T:31.3; A:27.2 |
| 152 | Co16_Stroke | C/t | T | 58.82T 41.18C | T:70.4; C:29.5; GAP:0.1; G:0.02 | 196 | 157 | 0,01469625 | C:47.9; T:35.4; A:16.7 |
| 16092 | Co40_Stroke | C/t | T | 82.35C 17.65C | T:98.7, C:1.2; Y:0,1; GAP:0.02 | 16 | 17 | 0,00159131 | A:41.7; T:33.3; C:20.8; GAP:4.2 |
| 16399 | Co62_Stroke | G/a | A | 83.33G 16.67A | A:97.4; G: 2.5; T: 0.02; C:0.02; GAP: 0.02; R:0.02 | 21 | 26 | 0,00243377 | T:39.6; A:31.3; C:16.7; G:6.3; GAP:6.3 |
| 146 | Ca120_MI | T/c | T | 63.64T 36.36C | T:81.4; C:18.4; A:0.1; GAP:0.1 | 121 | 109 | 0,01020313 | C:41.7; T:31.3; A:27.1 |
| 152 | Ca100_M | C/t | T | 52.94C 47.06T | T:70.4; C:29.5; GAP:0.1; G:0.02 | 196 | 157 | 0,01469625 | C:47.9; T:35.4; A:16.7 |
| 73 | Co6_MI | G/a | A | 76G 24A | G:80.8; A:19.1; GAP:0.1; C:0.02 | 12 | 11 | 0,00102967 | A:41.7; C:35.4; G:22.9 |
| 146 | Co56_MI | T/c | T | 66.67T 33.33C | T:81.4; C:18.4; A:0.1; GAP:0.1 | 121 | 109 | 0,01020313 | C:41.7; T:31.3; A:27.1 |
| 150 | Co17_MI | T/c | C | 86.20T 13.80C | C:88.2; T:11.7; G:0.1; GAP:0.1 | 74 | 63 | 0,00589722 | C:43.8; G:27.1; A:18.8; T:10.4 |
| 150 | Co123_MI | T/c | C | 82.76T 17.24C | C:88.2; T:11.7; G:0.1; GAP:0.1 | 74 | 63 | 0,00589722 | C:43.8; G:27.1; A:18.8; T:10.4 |
| 152 | Co27_MI | T/c | T | 58.82T 41.18C | T:70.4; C:29.5; GAP:0.1; G:0.02 | 196 | 157 | 0,01469625 | C:47.9; T:35.4; A:16.7 |
| 152 | Co184_MI | T/c | T | 52.38T 47.62C | T:70.4; C:29.5; GAP:0.1; G:0.02 | 196 | 157 | 0,01469625 | C:47.9; T:35.4; A:16.7 |
| 204 | Co156_MI | T/c | T | 83.33T 16.67C | T:93.4, C:6.5, A:0.1, GAP:0.1; Y: 0,02 | 44 | 43 | 0,00402509 | G:52.1; T:22.9; C:16.7; A:8.3 |
| 16092 | Co62_MI | C/t | T | 82.35C 17.65C | T:98.7, C:1.2; Y:0,1; GAP:0.02 | 16 | 17 | 0,00159131 | A:41.7; T:33.3; C:20.8; GAP:4.2 |
| 16093 | Co106_MI | C/t | T | 72.22C 27.78T | T:93.5, C:6.4; Y:0,1; GAP:0.02 | 55 | 79 | 0,00739493 | T:64.6; A:20.8; G:6.3; GAP:4.2; C:4.2 |
| 16129 | Co129_MI | G/a | A | 81.81G 18.19A | G:84.6; A: 14.9; C: 0.4; R:0.02; GAP: 0.02 | 93 | 86 | 0,00805017 | A:87.5; G:8.3; T:2.1; GAP:2.1 |
| 16399 | Co94_MI | G/a | A | 83.33G 16.67A | A:97.4; G: 2.5; T: 0.02; C:0.02; GAP: 0.02; R:0.02 | 21 | 26 | 0,00243377 | T:39.6; A:31.3; C:16.7; G:6.3; GAP:6.3 |

### Analysis of fixed and heteroplasmic mtDNA mutations with MI

MtDNA positions studied for MI cases and controls are available in the matrix of S2 Table and the frequencies of fixed mutations found are reported in Table 4. The m.72T>C, m.73A>G and m.16356T>C were more frequent in MI controls (12.3%, 49.3% and 3.3%, respectively) than cases (7.6%, 38.9% and 1.4%, respectively). When corrected for the effect of CV risk factors with significant differences between MI cases and control (hypertension and hypercholesterolemia [24]), significant association was observed in these tree mutations (m.72T>C: conditional logistic regression: p=0.001; OR= 0.041, 95% CI [0.006-0.290], m.73A>G: conditional logistic regression: p=0.009; OR= 0.009, 95% CI [0.307-0.843] and m.16356T>C: conditional logistic regression: p= 0.016; OR= 0.091, 95% CI [0.013-0.639]), emerging as a possible protective genetic factors for MI (Table 3).

**Table 4:** Complete results of myocardial infarction fixed mtDNA mutation analysed

|  | Position 72T>C | Position 73A>G | Position 16356T>C |
| --- | --- | --- | --- |
| Controls: n=211 (%) / Cases: n=211 (%) | 26 (12.3) / 16 (7.6) | 104 (49.3) / 82 (38.9) | 7 (3.32) / 3 (1.42) |
| Logistic Regression<sup>1</sup> |  |  |  |
| p-value | 0.001 | 0.009 | 0.016 |
| OR 95% CI | 0.041 0.006-0.290 | 0.509 0.307-0.843 | 0.091 0.013-0.639 |
| Stability analysis |  |  |  |
| CRS | T | A | T |
| Distribution in population database | T:97.4; C:2.4; G:0.1;<br>GAP:0.1 | G:80.8; A:19.1;<br>GAP:0.1; C:0.02 | T:97.5; C:2.5;<br>GAP:0.02 |
| No. Hits phylogeny (PhyloTree.org) | 9 | 12 | 15 |
| No. Hits Soares et al. 31 | 6 | 11 | 19 |
| Probability of mutation | 0,00056164 | 0,00102967 | 0,00177853 |
| Nucleotide Conservation Index (%) | T:77.1; GAP:8.3;<br>A:6.3; C:4.2; G:4.2 | A:41.7; C:35.4;<br>G:22.9 | T:79.2; C:12.5;<br>A:6.3; GAP:2.1 |

<sup>1</sup> Conditional regression model was performed considering significant covariates for MI (hypertension, hypercholesterolemia and mtDNA mutations)

In order to predict the impact of these mutations, several measures were calculated to analyze the stability of each position, and results are showed in Table 4. The results obtained revealed that m.72T>C, m.73A>G and m.16356T>C as a non-stable positions since they present a minimum of 9 hits in the phylogeny, a high probability of mutation, a high frequency of the variant in the population database (MAF >5%) for m.73A>G and low-frequency (MAF 1-5%) for m.72T>C and m.16356T>C, and a maximum nucleotide CI of 79% (Table 4). Using the proposed previously method to predict the impact of these three mutations on the stability of secondary structure of the mtDNA, it seems that m.72T>C, m.73A>G and m.16356T>C led to a folded structure with the same minimum free energy as the wild-type structure (rCRS), which means that these mutations do not condition the stability of the region.

Classification of the heteroplasmic positions between MI cases and control is available in Table 2. One hundred twenty-five MI cases (59.2%) and one hundred and twenty controls (56.8%) presented point and/or length heteroplasmy, being the length heteroplasmy located in the poly-C tract of the HVRII the most prevalent variant both MI cases (54.03%) and controls (48.34%). In this analysis, the frequency of point heteroplasmy was overrepresented in MI controls (n=11; 5.21%), differing significantly from cases (n=2; 0.94%) (p=0.022, McNemar’s test). This association remained significant (logistic regression: p=0.046; OR= 0.209, 95% CI [0.045-0.972]) even correcting for the effect of MI risk factors (hypertension and hypercholesterolemia [24]). These heteroplasmic positions involving nine different positions of the mtDNA: 73, 146, 150, 152, 204, 16092, 16093, 16129 and 16399.

The stability analysis to identify the impact of these point heteroplasmy is presented in Table 3. All of them were considered non-stable positions. As previously stated, these positions presented a minimum of 16 hits in the phylogeny, were located in hotspots positions, have a high frequency of the minor variant in the population database (min. height peaks 13.80%) and low CI at nucleotide level (max. 64.4%). No different trends were observed between stability of these positions in MI cases and controls (Table 3).

### Distribution of mtDNA mutations between haplogroups

Haplogroup assignment of all individuals analysed in this article was previously performed by Umbria et al. [24]. In Fig 2 are shown the distribution of m.16145G>A and m.16311T>C for stroke and m.72T>C, m.73A>G and m.16356T>C for MI between the mtDNA haplogroups. Unlike the rest of mutations, it appears that m.72T>C and m.16356T>C have a high association with the haplogroups HV0 and U, respectively (Fig 2). These associations ceases to exist when compared the number of individuals identify in this article that belonging the haplogrup HV0 and have m.72T>C with the total of HV0 cases and controls samples detected by Umbria et al. [24], shown that there are individuals belonging this haplogroup that do not have the m.72T>C. In the same line, note that the total number of individuals identify in this study that have m.16356T>C belongs to haplogroup U. Even though m.16356T>C define different subgroups of U (U2e3, U3a1c, U4 and U5b1), this result was obtained because in this analysis the haplogroup U included the rest of subhaplogrups of U detected. Otherwise, the frequencies obtained would be more divided. Hence, the analysis of the distribution of these mutations clearly demonstrated that they weren’t associated with any particular mtDNA haplogroup. Therefore, these mutations could act as independent risk factors for haplogroups.

**Fig 2.**
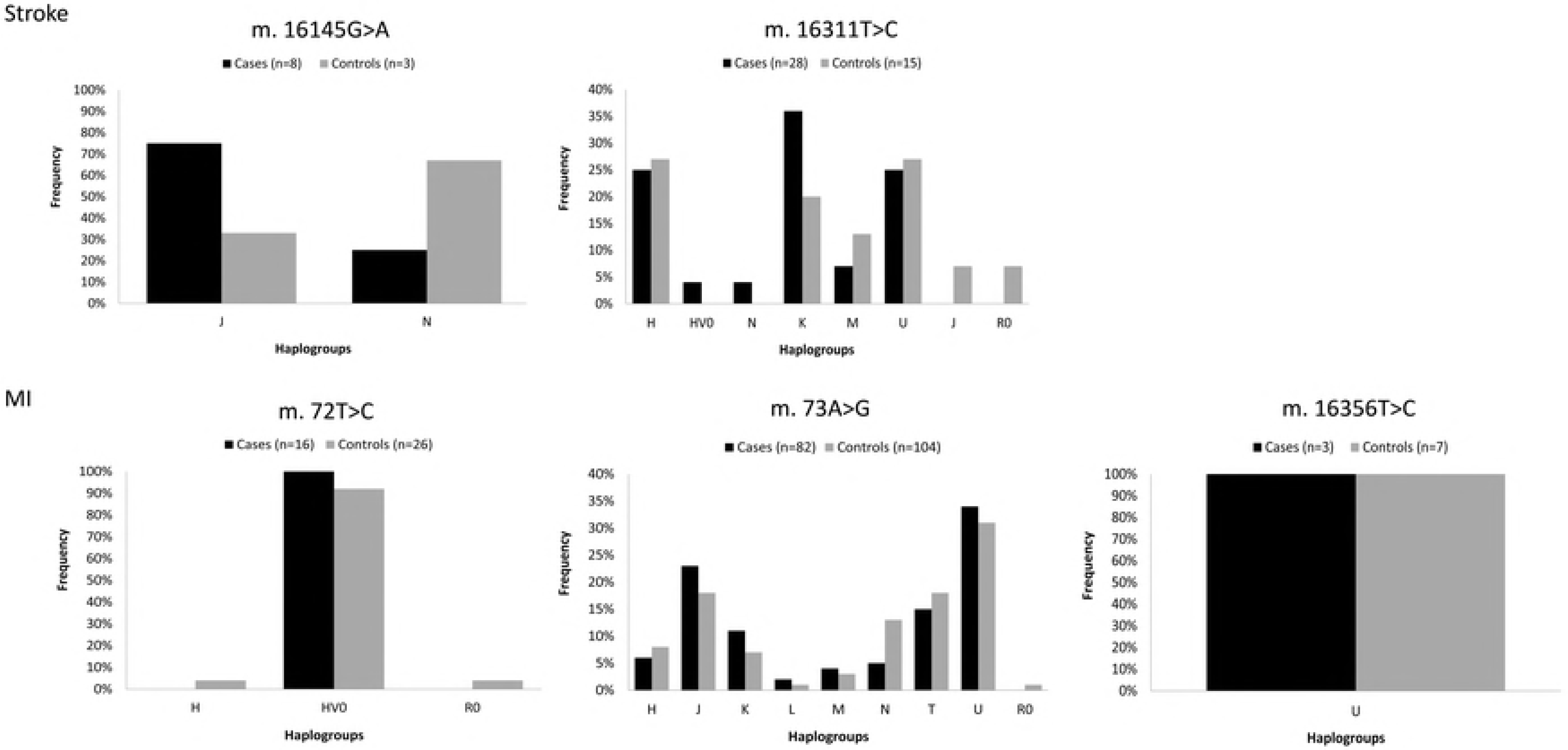
Distribution of fixed mutations between the mtDNA haplogruops

## Discussion

In western countries, where the burden of CVD is growing due to effect of CV risk factors, several studies have already shown the strongly relation of the genetic factors. However, little is know about the role of mtDNA CR mutations in development of stroke and MI [7–9,11,13,14,15,17].

An association of several mtDNA alterations (fixed and in heteroplasmy) in the two diseases have been detected in the present study. As regards fixed mtDNA mutations, the set of mutations in stroke and MI cases was compared to controls and significant differences were found in the two diseases, reporting the m.16145G>A and m.16311T>C as a potential genetic risk factors for stroke, and m.72T>C, m.73A>G and m.16356T>C as a possible beneficial genetic factors for MI. It has been previously described that the CR mutations can be associated across multiple disease, and that the same variant could had opposite effect (increase or decrease the risk) for two different disease [10]. This finding would support our original hypothesis about the consequences that can affect the mtDNA mutations in the CR depending on the disease.

The CR variants do not act directly on the ETC affecting mitochondria bioenergetics or ROS generation; they may have an important effect on the genotype by altering mtDNA gene expression [18,19]. The CR contains the main regulatory sequences for replication initiation and transcription [3]. Unfortunately, the potential role of the remaining areas of the CR is still unknown even considering the exceptional economy of organization of the mtDNA [26].

Transitions 16145G>A and 16311T>C seem to have a pathogenic role in stroke. The analysis of distribution of these mutations clearly showed that they are located in many different haplogroups and consequently these mutations act as haplogroup-independent risk factors. The m.16145G>A is located between MT-TAS sequence (nt. 16157-16172) and MT-TAS2 sequence (nt. 16081-16138). According to the classic strand-asynchronous mechanism, recent studies demonstrated that the 5’end of the D-loop is capable of forming secondary structures [27], which act as a recognition site to molecules involved in the premature arrest of H strand elongation [28]. The biological importance of this region was confirmed by Brandon et al. [29] who also observed multiple tumor specific mutations in the pre-TAS region. These observations suggest that mutations arising near to this conserved motive might be responsible of the alterations in mtDNA replication and transcription. In the same line, m.16311T>C has been found to be significantly associated with certain types of cancer [30–32]. This mutation was previously described by Chen et al. [30] in patients with prostate cancer and also has been reported in colorectal cancer [31] and more recently in acute myeloid leukemia [32]. This mutation is located between the control elements Mt5 sequence (nt. 16194-16208) and the Mt3l sequence (nt. 16499-16506). In this case, our results showed that m.16311T>C may implies a reduction in the stability of secondary structure of this region, which would affect in the binding grade to mtDNA transcription factors, ultimately affecting on the intensity of transcription regulation [33]. In both cases, these findings strongly suggest that mtDNA CR dysfunction may cause a decrease on the mtDNA copy number, which could affect the efficiency of ETC, lowering the ATP:ADP ratio and increasing ROS production [18,19], contributing in stroke development.

Concerning MI, our results showed that m.72T>C, m.73A>G and m.16356T>C act as a beneficial factor for MI. Although a high percentage of individuals with these mutations belonged to the haplogroups HV0, H or U, which have been shown to may have higher oxidative damage [9,24,34,35], the distribution of positions 72, 73 and 16356 in our samples was independent of these haplogroups. Since the role of the mitochondrial genome in CVD susceptibility remains uncertain, it is difficult to explain how these mutations can decrease or counteract the progression of MI. Although m.72T>C, m.73A>G and m.16356T>C have been previously related to certain types of cancer [36], many studies consider that they are recurrent variants common in humans [37].

Even though the most deleterious mutations are removed by natural selection, a wide range of milder bioenergetic alterations are introduced in certain populations [38]. Some of these variants as m.72T>C, m.73A>G and m.16356T>C could be advantageous and seen as way to facilitate survival in specific environments. In contrary, other mutations as m.16145G>A and m.16311T>C escape of intraovarian selection and could cause significant mitochondrial defects and stroke develop. Much of the progress in linkage disequilibrium mapping of complex diseases has been made using the major assumptions of the CDCV hypothesis, that is, that common alleles cause common diseases. After found positive associations with common alleles (e.g., those found by Umbria et al. [24]), it was necessary replicated the results and then look for rarer variants, with potentially greater penetrance. However, all the mutations analysed in the present study had a minor allele variant >5% (common variant) or between 1-5% (low-frequency). Although the common variants often are associated with OR of only between 1.2 and 1.5 [39], our results showed high effect size for pathogenic variants, with OR values of 2.4-4.4, and similarly, high protective effect size of variants found at higher frequency in controls (ORs <0.5) relative to cases.

Our findings also showed a significant increase of point heteroplasmy in MI controls in comparison to cases. This result is contrary to expectations, because the presence of heteroplasmy has been commonly associated with aging and degenerative disease, due to a decline in mitochondrial function in both these processes [40]. Our registered heteroplasmic positions (16399, 16129, 16093, 16092, 73, 146, 150, 152, and 204) were located in hotspots positions of the hypervariable segments. Recent evidences demonstrate that an important fraction of mutations detected in heteroplasmy are germinal or originated in very early stages of the development [41]. Moreover, it is probable that germinal heteroplasmy has a beneficial or risk effect, and our results revealed that the higher number of point heteroplasmy were overrepresented in MI controls individuals. This fact, is not surprise because some heteroplasmic positions detected, as m.73G>A, has been linked in this study as a possible beneficial genetic factors for MI and also it has been suggested that other heteroplasmic positions, such as 146T>C, 150C>T or 152T>C may increase longevity [42].

Many studies have shown that heteroplasmic variants without apparent functional consequences are observed in apparently healthy individuals [43,44,45,46]. In the present study, the frequency of MI controls with point heteroplasmy in the CR (5.2%; 95% CI [0.045-0.972]) are slightly higher than those reported by Santos el at. [45] (3.81%; 95% CI [0.166−0.737]), but less than described by Ramos et al. [46] (7.9%; 95% CI [0.041− 0.149]), demonstrating that heteroplasmy occur with appreciable frequency in the general population [43]. This idea is even more reinforced in front of the present data, since the high representation of point heteroplasmies detected were located in non-stable positions and by the fact that no significant differences were found between the frequencies of point heteroplasmy in stroke cases and controls in the present study.

In conclusion, our finding indicates the possible role of mtDNA CR mutations in the pathogenesis of stroke and MI. Our results may provide better understanding of the cellular mechanism by which mtDNA variants contribute to CVD, and endorse the importance of including this regulatory region of the mtDNA in genetic association studies.

## Material and methods

### Subjects

In this study, data from 730 subjects (154 individuals with stroke history, 211 individuals with MI history and their corresponding control individuals -matched for age, gender and geographic origin-, were used. Samples come from the Cardiovascular Disease Risk Study of Castile and Leon [47], whose design and analysis have already been described by Umbria et al. [24]. For each individual, we also obtained information about history of hypertension (≥140/90 mmHg), history of diabetes, history of hypercholesterolemia (>200 mg/dl), cigarette consumption (smokers, former smokers and non-smokers), presence of overweight or obesity (body mass index ≥25 kg/m2), presence of high abdominal perimeter in risk range (risk: ≥80 cm for women and ≥94 cm for men) and presence of high levels of triglycerides (≥170 mg/dl).

### MtDNA sequence analysis and heteroplasmy authentication

The mtDNA sequences used in the present study were previously obtained by Umbria et al. [25] although, they were strictly used to classify samples into mtDNA haplogroups. In the present study, sequences were reassessed and analysed at the nucleotide level to identify not only fixed mutations but also mutations in heteroplasmy. The alignment in relation to the to the revised Cambridge Reference Sequence (rCRS) [26] and the heteroplasmy detection were performed using the SeqScape 2.5 software (Applied Byosistems, Foster City, USA) considering a value of 5% in the Mixed Base Identification option. Only sequences with satisfactory peak intensity and without background/noise were considered. In this context, some samples were amplified and sequenced several times (using the same methodology described in Umbria et al. [25]) to obtain accurate sequences to heteroplasmy detection. Moreover, additional analyses were performed in order to authenticate heteroplasmies.

The authentication of mtDNA heteroplasmy was performed following a similar strategy to that used by Santos et al. [45,49].

1. PCR amplification and sequencing of the control region of the mtDNA.
2. To authenticate the results for samples presenting heteroplasmy in step 1, a second PCR amplification and sequencing were performed.
3. In addition, to exclude a possible contamination of the samples, an analysis of Short Tandem Repeat (STR) DNA profiling was carried out employing AmpFlSTR^®^ Identifiler^®^ PCR Amplification Kit (Applied Biosystems, Foster City, USA) following the manufacturer’s protocol.

Thus, point heteroplasmic positions were accepted if they appeared in all the validation steps and no evidence of sample contamination was detected.

Levels of heteroplasmy were determined using the height of peaks in the electropherograms [45]. To calculate the average heteroplasmic levels, the results obtained for at least two sequence reads of each heteroplasmic position were used.

### Data Analysis

#### Statistical analyses

To compare differences in the CR profile between cases and controls in both stroke and MI, all fixed and heteroplasmic mtDNA mutations were compiled into a matrix considering the cases and controls analysed for each disease.

The set of fixed mtDNA mutations detected in cases and controls (present in a minimum of two individuals) were compared using McNemar’s test. To adjust the association analysis for the potential confounding effect of CV risk factors detected previously in Umbria et al. [24], a conditional logistic regression analysis (forward stepwise model) was applied. Hypercholesterolemia was considered a CV risk factor with a potential confounding effect for stroke, while both hypertension and hypercholesterolemia were considered for MI samples. Therefore, Odds Ratios (ORs) and their 95% Confidence Intervals (CIs) were calculated adjusting for the effect of these risk factors in each disease.

McNemar’s test was applied to compare the presence or absence of point (PH) and length (LH) heteroplasmy, and a logistic regression analysis was used to correct for the effect of CV risk factors above mentioned [24].

Finally, the mtDNA mutations were analysed to evaluated if they were in positions defined by haplogroups (previously examined in Umbria et al. [24]), or acts as an independent genetic factor.

Statistical analyses were performed using IBM SPSS ver. 22.0 (SPSS Inc.). All differences were considered significant at p<0.05.

### Hits in the phylogeny, population database and Conservation Index (CI)

The stability of fixed mtDNA mutations and point heteroplasmic position were analysed as previously detailed by Ramos et al. [46]. The number of hits in the phylogeny for each position was compiled from the updated mtDNA phylogeny – mit. Tree build 17 [50] – and from Soares et al. [25]. From these data, it has been possible to calculate the probability of mutation as the ratio between the observed and the total number of hits. An mtDNA position was considered a hotspot if the mutation probability was ten time higher that the expected mean value. In order to calculate the frequency of each variant for a particular nucleotide position, a database of 3880 mtDNA complete sequences was used. Sequences were aligned using Clustal W and formatted for further frequency analyses using the SPSS software. The nucleotide conservation index (NCI) was estimated only across reference sequences of different primate species (for the list of species and accession numbers see S1 Table). Sequences were analyzed using the same method previously mentioned [46].

### Structure prediction

Secondary structures were performed to understand the structural impact of different variants found. The secondary structures for each position was generated from sequences (A-M) identifies by Pereira et al. [27]. All sequences were submitted to the RNAfold web server (http://rna.tbi.univie.ac.at/cgi-bin/RNAWebSuite/RNAfold.cgi) using default parameters for DNA secondary structures calculations. The minimum free energy prediction and base pair probabilities were used to estimate the implication in the molecule.

## Acknowledgements

This work was supported by MINECO (project: CGL2014-53781-r) and by Generalitat de Catalunya (Ref. 2017 SGR 1630).

Postdoctoral fellowship SFRH/BPD/105660/2015 (AR) was supported by Fundação para a Ciência e a Tecnologia (FCT).

## Supporting information

**S1 Table. List of accession number and primates used for the conservation index estimations at nucleotide level for D-loop.**

**S2 Table. Mutation report of mtDNA control region of 154 individuals with a history of stroke and 154 paired controls (Stroke case-control) and 211 individuals with a history of MI and 211 paired controls (MI case-control).**

